# Modeling Microbial Regulatory Feedback in Organic Matter Decomposition Identifies Copiotrophic Traits as Key Drivers of Positive Priming

**DOI:** 10.1101/2024.08.11.607483

**Authors:** Firnaaz Ahamed, James C. Stegen, Emily B. Graham, Timothy D. Scheibe, Hyun-Seob Song

## Abstract

Microbial decomposition of complex soil organic matter (OM) is often regulated by labile organic carbon inputs, a phenomenon known as priming, which plays a critical role in belowground biogeochemical cycling. However, the strength and direction of microbial priming of soil OM pools varies significantly across ecosystems. A generalizable mechanistic framework explaining the factors that lead to accelerated (positive priming) or impeded (negative priming) rates of OM decomposition is still lacking. In this work, we conceptualize priming as a microbial feedback loop that optimizes the costs and benefits of maximizing growth rate, specifically, the cost of exoenzyme synthesis for decomposing complex OM versus the benefit of energy acquisition from labile OM. We examined the impacts of microbial growth traits and interactions on priming by employing a cybernetic modelling approach, which predicts complex microbial growth patterns by accounting for dynamic metabolic regulations. We simulated microbial priming across ecological community configurations composed of degraders and non-degraders with either oligotrophic or copiotrophic growth traits, resulting in seven combinations that included both single functional groups (degraders with either growth trait) and binary functional groups (combinations of degraders and non-degraders, or degraders only, with differing or common traits). Configurations with only non-degraders were excluded, as they are irrelevant for studying priming in OM decomposition. Monte Carlo simulations for these scenarios revealed: (1) positive priming is prevalent, while negative priming occurs sporadically under specific parameter settings; (2) positive priming is more frequently observed in microbial systems with copiotrophic degraders than those with oligotrophic degraders; (3) the presence of copiotrophic non-degraders suppresses positive priming, whereas the presence of oligotrophic non-degraders promotes it; and (4) the temporal dynamics of priming is also influenced by microbial growth traits and interactions. These findings highlight the driving role of microbial functional traits and interactions in priming. Most strikingly, our simulations predicted a dramatic positive priming effect triggered by the addition of a small amount (i.e., less than 10%) of labile OM, with no notable changes observed beyond this point. As we used a generalized microbial model, we hypothesize that our findings may reflect common features of OM priming across diverse microbial systems and environments. Overall, this work, combining new theories and models, significantly enhances our understanding of priming by providing model-generated and empirically testable hypotheses on the mechanisms governing it.

## 1. Introduction

Priming is a phenomenon in which the introduction of chemically labile organic matter (OM) into an environment significantly influences microbial activity, leading to either accelerated (positive) or suppressed (negative) decomposition of more recalcitrant OM (i.e., chemically complex and/or stabilized) and other biogeochemical pools (e.g., minerals, nutrients) (Bingeman et al., 1953; Kuzyakov et al., 2000). The occurrence of priming has been frequently debated in the literature. For example, labile OM in the form of plant litter leachate and root exudates accelerate microbial decomposition of soil OM, which facilitates nutrient and energy exchange between soil microbial communities and plants in the rhizosphere (Fontaine et al., 2003; Soong et al., 2020). Priming equally manifests in other ecological domains, such as sediments in hyporheic zones of rivers and deep ocean seabed, which influence nutrient and energy cycles (Arrieta et al., 2015; Graham et al., 2017; Stegen et al., 2018). However, positive priming can also have adverse implications, as it stimulates increased microbial respiration, leading to heightened carbon dioxide emissions from the decomposition of soil OM rich in carbon (Nottingham et al., 2009; King, 2011). Conversely, negative priming aids carbon sequestration efforts by preserving soil carbon from mineralization (Guenet et al., 2018; Liang et al., 2023).

Despite its potential to exert long-term effects on ecosystem dynamics (King, 2011), our understanding of the fundamental mechanisms and key factors governing priming remains limited, posing challenges in accurately predicting biogeochemical dynamics. Priming can be influenced by biotic and abiotic factors, including microbial traits (Nottingham et al., 2009; Fontaine et al., 2011), environmental constraints such as carbon limitations (Graham et al., 2017; Soong et al., 2020), nutrient limitations (Fontaine et al., 2011; Hicks et al., 2019; Feng et al., 2021), or combinations thereof. Although experimental evidence suggests that microbial traits and interactions can play pivotal roles in priming (Brant et al., 2006; Garcia-Pausas and Paterson, 2011; Yu et al., 2018; Hicks et al., 2019), extensive microbial diversity in natural ecosystems poses obstacles to completely resolving their effects on OM decomposition (Trivedi et al., 2013).

We hypothesize that a key mechanism driving priming lies in the responses of microbial growth to the availability of labile OM. When new labile substrates are introduced, microbes face metabolic decisions that affect the allocation of cellular resources toward growth and exoenzyme production. This regulatory response directly determines the balance between energy acquisition from labile OM and the investment in decomposing complex OM, ultimately influencing whether positive or negative priming is observed. Importantly, such resource allocation strategies are governed by fundamental microbial traits that, at a coarse-grained level, can be categorized into two broad groups: copiotrophs (fast-growing microbes adapted to nutrient-rich environments) and oligotrophs (slow-growing microbes specialized for nutrient-poor conditions). Framing microbial priming through the lens of these growth strategies of OM degraders provides a tractable approach to predict priming responses across diverse systems.

In addition to degraders, non-degrading microbes that exploit labile OM produced through exoenzyme-mediated breakdown without contributing to enzyme production can further alter substrate dynamics and priming outcomes. These non-degraders can outcompete degraders for labile substrates, reducing the incentive for exoenzyme investment and potentially suppressing OM decomposition. Therefore, microbial interactions involving degraders and non-degraders are likely critical components of priming dynamics, yet remain underexplored in existing models (Momeni et al., 2013; Trivedi et al., 2013; Song et al., 2014).

Given that microbial regulation is inherently dynamic and context-dependent, it is also important to understand how priming patterns evolve over time and whether there are discernible temporal patterns or phases that reveal the underlying factors. Indeed, priming effects are time-sensitive phenomena that depend on the period over which they are evaluated (Zhang et al., 2017), influencing both their magnitude and direction. Therefore, exploring the time dependency of priming is of critical importance in quantifying priming effects across systems with different timescales.

To achieve a mechanistic understanding of priming, it is crucial to simultaneously consider microbial growth traits, microbial interactions, dynamic regulated synthesis of exoenzymes, and the role of non-degraders (Fontaine et al., 2003; Brant et al., 2006; Garcia-Pausas and Paterson, 2011; Di Lonardo et al., 2017; Yu et al., 2018; Hicks et al., 2019). The limited consideration of these microbial processes in most studies hinders the elucidation of priming mechanisms (Nottingham et al., 2009; Fontaine et al., 2011; Trivedi et al., 2013). Moreover, experimental approaches quantifying priming effects through changes in respiration rates as proxies for OM decomposition also face limitations due to the associated complexities with distinguishing real priming from apparent changes (e.g., microbial biomass turnover). In essence, there are currently no generalizable theories that fully explain the diverse patterns of priming effects, highlighting the need for mechanistic models capable of predicting both positive and negative priming within a unified framework.

In this regard, cybernetic modeling is of particular interest due to its unique capability of predicting complex microbial growth patterns in dynamically varying environments by accounting for metabolic regulation (Ramkrishna and Song, 2018). Instead of accounting for all molecular details of microbial regulation that are generally unknown, cybernetic models provide rational descriptions of regulation based on cybernetic laws derived from an optimal control theory. Cybernetic control laws dictate which enzymes microbes synthesize and activate to maximize their growth rates under given environmental conditions (Young and Ramkrishna, 2007), balancing the cost of enzyme synthesis for complex OM decomposition against the energy gained from labile OM, following an economic return-on-investment concept. Therefore, cybernetic modeling can serve as an ideal tool for understanding priming because it accounts for microbial regulation through a feedback loop comprising three key processes: enzyme synthesis, complex OM decomposition, and microbial growth on labile OM.

In this work, we propose a novel theoretical framework that employs cybernetic modeling to predict microbial responses to the addition of labile OM and their impacts on the decomposition rates of chemically complex OM. Towards identifying key factors governing priming effects, we evaluate several questions: (1) how growth traits of microorganisms degrading complex OM drive positive and negative priming, (2) how the presence of non-degraders affect priming through interactions with degraders, (3) how the temporal patterns of priming are affected by these factors, and (4) whether there are any common features of priming predicted by theoretical models across different combinations of microbial traits. We modeled a targeted set of microbial group combinations needed to evaluate these questions. These combinations had both single and binary functional groups of degraders and non-degraders, each with either copiotrophic or oligotrophic growth traits. Grounded in a unified framework, we explored how microbial growth traits and interactions influence priming rates. To draw unbiased conclusions, we conducted Monte Carlo simulations by assigning random parameter values to each model, rather than calibrating parameters with specific values that might be relevant to one system but not to others. The comprehensive simulations generated by this method provided new insights into each of the key questions, revealing several critical aspects of priming.

## 2. Methods

### 2.1 Development of a general theoretical model to elucidate the impact of microbial growth traits and interactions on priming effects

Our modeling framework considers the mixing of relative amounts of complex and labile OM (**Fig. 1A**) as the chemical parameter governing the occurrence of positive or negative priming, whereas the biological parameter is represented by three fundamental microbial processes: synthesis of extracellular enzymes (i.e., exoenzymes), decomposition of complex OM, and microbial growth on labile OM (**Fig. 1B**). These processes interact with each other through a closed feedback loop (Ahamed et al., 2021). Microorganisms synthesize exoenzymes to break down complex OM, producing labile OM that can be readily assimilated to support cell growth. In unprimed conditions, this feedback loop is inactive or slow, but when perturbed by the addition of exogenous labile OM, microorganisms gain additional energy to support growth and synthesize further exoenzymes, potentially facilitating positive priming by accelerating the decomposition of complex OM. The feedback loop can be suppressed if the energy return from degrading complex OM is not favorable to support cell growth, potentially leading to zero or even negative priming. The framework further expounds the possibility that the combined microbial cost-benefit regulations and interactions between microbial groups with distinct growth traits lead to various patterns of priming effects. The following microbial group combinations are the primary focus of this study, as the subset of possible combinations that most directly address our stated science questions: single functional groups (**Fig. 1C**) consisting of (1) a degrader with copiotrophic growth traits (CDG), and (2) a degrader with oligotrophic growth traits (ODG); binary consortia (**Fig. 1D**) consisting of (3) a pair of CDG and oligotrophic non-degrader (OND), and (4) a pair of copiotrophic non-degrader (CND) and ODG.

**Figure 1.**
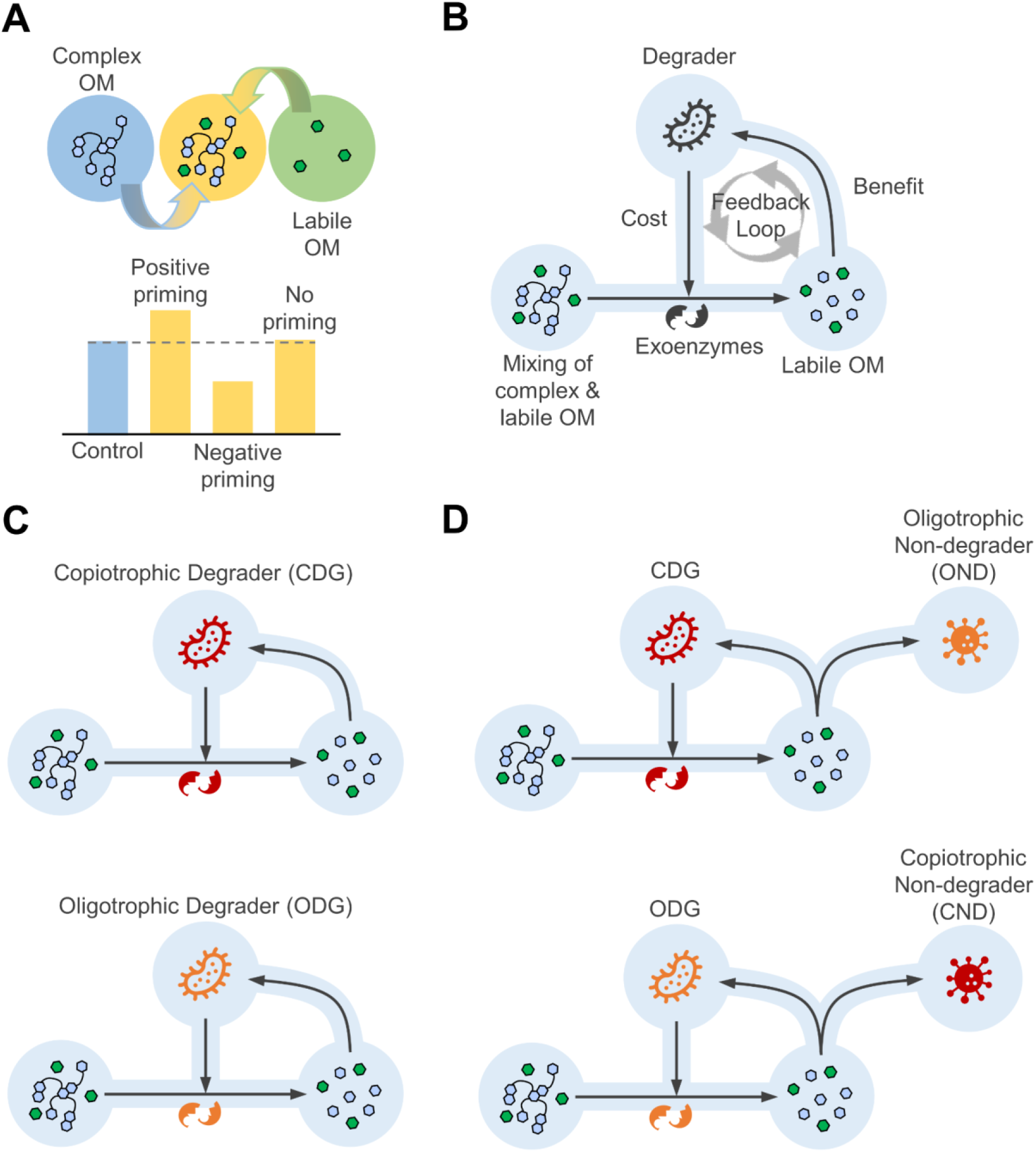
An illustration of a conceptual model developed to simulate priming effects driven by **(A)** chemical factors, specifically the mixing of complex and labile OM, which results in varied levels of complex OM decomposition rates, leading to positive, negative, or no priming. This process is governed by **(B)** microbial factors, i.e., dynamic cellular regulation through a closed feedback loop, which manages resources to balance the costs of exoenzyme synthesis for degrading complex OM and the benefits of assimilating produced labile OM for cell growth. Considering the influence of distinct microbial growth traits (oligotrophs and copiotrophs) and their interactions on priming, four major priming models are obtained depending on how copiotrophic and oligotrophic growth traits are assigned in **(C)** single functional groups with degraders alone, and **(D)** binary consortia with degraders and non-degraders.

### 2.2 Model equations

For generality, we provide the model equations only for a binary consortium composed of microbial species 𝑖 and 𝑗 representing mixed growth traits because the single functional group models are readily derivable from the consortium model by removing one of the species. The growth of species 𝑖 in the consortium in a uniformly mixed batch environment is represented as:

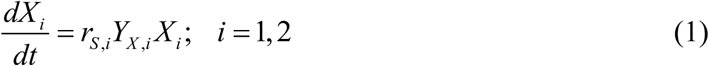

where the subscript 𝑖 denotes the 𝑖^th^ species, 𝑋_𝑖_, 𝑌_𝑋,𝑖_, and 𝑟_𝑆,𝑖_ represent biomass concentration, biomass yield, and specific growth rate, respectively. The mass balance of the labile OM is given by:

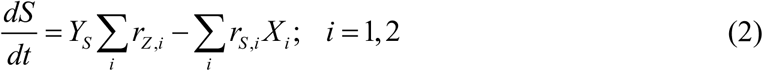

where 𝑌_𝑆_ is the yield of labile OM from the degradation of complex OM, 𝑟_𝑍,𝑖_ is the rate of degradation of complex OM by species 𝑖, and 𝑟_𝑆,𝑖_ is the specific uptake rate of labile OM in species 𝑖. Accordingly, the degradation of complex OM is given as:

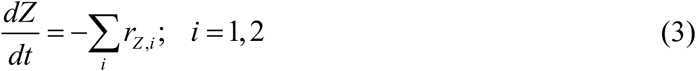

The degradation of complex OM and the assimilation of labile OM is regulated by exoenzymes and endoenzymes, respectively:

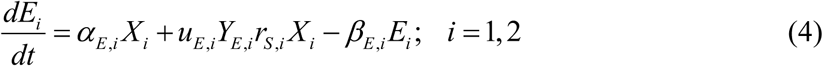

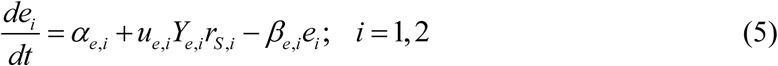

Here, 𝛼_𝐸,𝑖_ and 𝛼_𝑒,𝑖_ are the constitutive (basal) enzyme synthesis rates, 𝑌_𝐸,𝑖_ and 𝑌_𝑒,𝑖_ are the enzyme yields, and 𝛽_𝐸,𝑖_ and 𝛽_𝑒,𝑖_ are the enzyme decay rates. Then, the specific uptake rate of labile OM and the degradation rate of complex OM are given in the form of Michaelis-Menten kinetics:

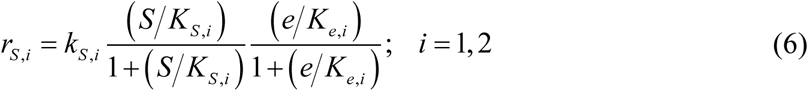

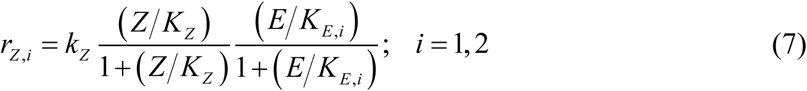

Here, 𝑘_𝑆,𝑖_ and 𝑘_𝑍_ are the maximum rates of labile OM uptake and complex OM decomposition, respectively, and 𝐾_𝑆,𝑖_, 𝐾_𝑍_, 𝐾_𝑒,𝑖_ and 𝐾_𝐸,𝑖_ are their associated saturation constants.

In single functional group models, we consider that microorganisms can degrade complex OM (i.e., degraders). In binary consortium models, we consider one species to be a degrader and the other species to be a non-degrader.

### 2.3 Parameter assignment and simulation settings

Variability in model parameter values leads to drastically different priming outcomes. Therefore, predicting priming effects is challenging even with known microbial growth traits and environmental conditions. To address this, we generate model results using Monte Carlo simulations by assigning random values to model parameters across different environmental conditions except the kinetic parameters associated with microbial growth. That is, to reduce the complexity of analysis, we fixed the labile OM uptake kinetic parameters (𝑘_𝑆,𝑖_, 𝐾_𝑆,𝑖_) to distinguish the growth traits between oligotrophs and copiotrophs (see **Supplementary Text** and **Supplementary Fig. S1**). Similarly, we fixed the exoenzyme synthesis kinetics to differentiate between degraders (𝑑𝐸_𝑖_⁄𝑑𝑡 ≠ 0) and non-degraders (𝑑𝐸_𝑖_⁄𝑑𝑡 = 0) based on their ability to synthesize exoenzymes. Combination of the role of microorganisms (as degraders or non-degraders) with copiotrophic and oligotrophic growth traits based on this pre-chosen parameter setting generates the four major microbial groups considered in the simulations: copiotrophic degrader (CDG), copiotrophic non-degrader (CND), oligotrophic degrader (ODG), and oligotrophic non-degrader (OND). To ensure a fair comparison between different test cases, the initial microbial population densities are made equal between the two microbial groups in the binary consortia. The same population densities are also used in the single functional group models, meaning the total initial biomass in these models is half that of the consortium models.

Importantly, the complex OM degradation kinetic parameters (𝑘_𝑍_, 𝐾_𝑍_, 𝑌_𝑆_) were also fixed to ensure that this study focuses solely on the variability in the growth traits of the microbes affecting the priming outcome, rather than the variability in complex OM properties. To represent the degradation of complex OM and the production of labile OM, 𝑌_𝑆_ must be greater than 1 (synonymous with the degradation of long-chain polymers into shorter oligomers or monomers) but using different values would only adjust the magnitude of priming without altering the qualitative trends. In other words, only the exoenzyme and endoenzyme synthesis kinetic parameters are randomized in the Monte Carlo simulations. By stochastically assigning values to these model parameters within prescribed ranges using a uniform distribution, we investigated the combined effects of microbial cost-benefit regulations and interactions between microbial groups with distinct growth traits on priming outcomes. To vary the environmental conditions to manifest different priming patterns, we varied the mixing fraction of labile OM with complex OM from 0 to 1, representing a continuum from pure complex OM to pure labile OM, respectively. In all cases, the Monte Carlo simulation results were based on no less than 200 runs for each mixture. The values of all model parameters, including those that were fixed and the bounds for randomized parameters, are provided in **Supplementary Table S1**.

### 2.4 Representation of microbial regulation using cybernetic modeling

The cost-benefit regulation of exoenzyme synthesis, and energy acquisition from microbial growth achieved from the assimilation of labile OM through the synthesis of exo- and endoenzymes in Eqs. (4) and (5), are governed by cybernetic variables 𝑢_𝐸,𝑖_ and 𝑢_𝑒,𝑖_, respectively, where 𝑢_𝐸,𝑖_ + 𝑢_𝑒,𝑖_ = 1. We employ generalized cybernetic laws based on optimal control systems, enabling the representation of microorganisms’ evolutionary strategy to regulate their metabolism in the interest of future returns over a finite time horizon (Young and Ramkrishna, 2007). This modeling capability is particularly crucial in complex substrate environments, where the investment of invaluable cellular resources for synthesizing complex OM-degrading exoenzymes does not yield immediate returns in terms of cell growth but is necessary for future growth when degraded products (labile OM) become available for assimilation. The detailed formulation of optimal control-based generalized cybernetic regulation is given in Young and Ramkrishna (2007), but briefly:

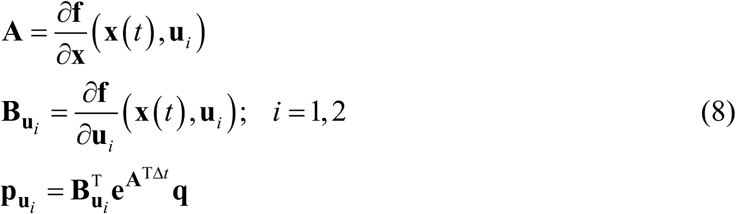

Here, 𝐟 is the system of ordinary differential equations in Eqs. (1) – (5), 𝐱 is the state vector of the system, 𝑢 = [𝑢 , 𝑢 ]^T^, and 𝐪 is the vector representing the metabolic objective of the system, which in this work is the cell growth rate of the respective microbial groups, given by Eq. (1). The future finite time horizon across which the returns are evaluated is computed based on the minimum time scale of the system, i.e.,

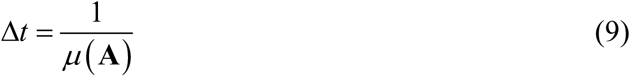

where 𝜇(𝐀) is the maximum eigenvalue of A. Subsequently, the cybernetic variables are evaluated using matching law over the return-on-investment of finite cellular resources:

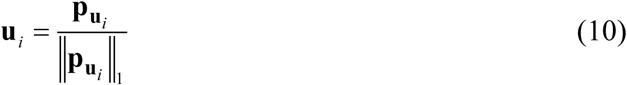

### 2.5 Metrics for quantifying model-estimated priming effects

Priming effects refer to the change in the decomposition rate of complex OM when labile OM is added to the system. Experimental studies of priming effects often face limitations as they rely on changes in microbial respiration rates as a proxy for OM decomposition, which may not be ideal. However, in our modeling framework, we can directly quantify the overall relative priming effects by comparing the amounts of degraded complex OM in perturbed and unperturbed systems over a specified period:

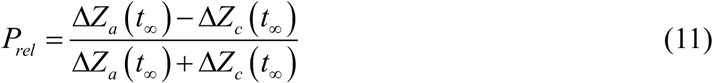

Here, Δ𝑍_a_ and Δ𝑍_c_ are the amended (treated with exogenous labile OM) and control amount of degraded complex OM evaluated at a time 𝑡, respectively. Here, the overall relative priming effect is normalized such that the maximum positive priming (𝑃_rel_ = +1) is realized when Δ𝑍_a_ ≫ Δ𝑍_c_, maximum negative priming (𝑃_rel_ = −1) is attained when no amended complex OM is degraded (Δ𝑍_a_ = 0), and no priming effect (𝑃_rel_ = 0) is observed when Δ𝑍_a_ = Δ𝑍_c_. For special cases where Δ𝑍_a_ = Δ𝑍_c_ = 0, we assign no priming effect, i.e., 𝑃_rel_ = 0. To compare the priming effects consistently across test cases with different complex OM decomposition dynamics and timescales following the stochastic modeling, we define 𝑡_∞_ as the moment when the complex OM is depleted to within a fixed tolerance:

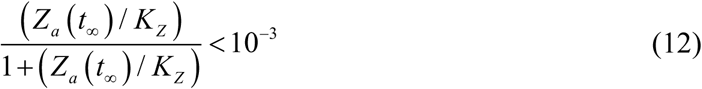

The threshold above is low enough that it will no longer drive priming in the system unless external disturbances are re-introduced. Subsequently, the instantaneous relative priming effects are represented as:

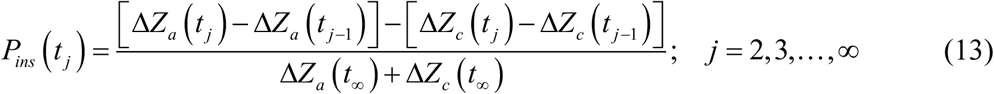

where the overall relative priming effects and the instantaneous relative priming effects are associated such that:

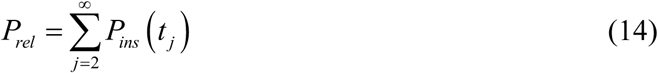

## 3 Results

Using the microbial priming model described in the previous section, we simulated seven cases representing distinct functional traits of degraders and non-degraders. To maintain clarity, we focus in the following sections on four major configurations, including CDG, ODG, CDG–OND, and ODG–CND, illustrated in **Figs. 1C** and **D**. Additional microbial group combinations, including ODG–OND, CDG–CND, and CDG–ODG, were also analyzed to encompass all possible pairwise configurations (**Supplementary Text** and **Fig. S2**). A summary of priming outcomes across all test cases is provided in **Supplementary Table S2**.

### 3.1 Copiotrophic growth shows modestly stronger positive priming than oligotrophic growth

We examined how priming is affected by the growth traits of degraders, i.e., copiotrophs and oligotrophs, using the single functional group models illustrated in **Fig. 1C**. We compared the levels of overall relative priming (𝑃_𝑟𝑒𝑙_) and biomass concentrations (𝑋) as a function of mixing fractions of labile OM (𝜈) with complex OM, between the CDG and ODG models in **Fig. 2**. Positive and negative values of 𝑃_𝑟𝑒𝑙_ represent the occurrence of positive and negative priming, respectively.

**Figure 2.**
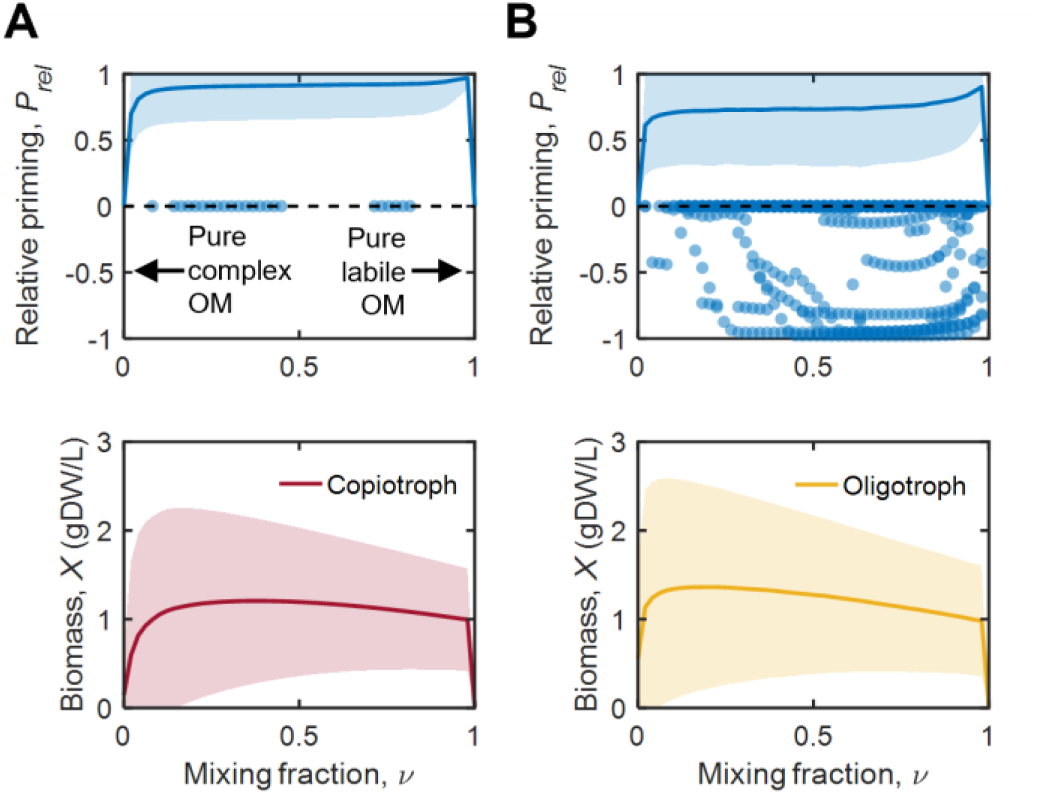
Overall relative priming effects and microbial population levels at various mixing compositions of complex and exogenous labile OM, evaluated for single functional group models of **(A)** copiotrophic degraders, and **(B)** oligotrophic degraders. The lines and shaded regions are averaged results and standard deviations of Monte Carlo simulations, respectively, while the markers are individual runs with negative overall relative priming effects.

Both models (top panels of **Fig. 2**) predicted that the average 𝑃_𝑟𝑒𝑙_ values were zero at unmixed conditions as expected, i.e., 𝜈 = 0 with pure complex OM or 𝜈 = 1 with pure labile OM. The values drastically increased around 𝜈 ≈ 0.1, and then remained almost constant until reaching a maximum around 𝜈 ≈ 0.97. The CDG model predicted the prevalence of positive priming, while also showing instances where 𝑃_𝑟𝑒𝑙_ values were zero, depending on the parameter sets randomly assigned in individual Monte Carlo simulations (top panel of **Fig. 2A**). The ODG model also predicted the dominance of positive priming but showed many instances where individual 𝑃_𝑟𝑒𝑙_ values were negative, resulting in lower average 𝑃_𝑟𝑒𝑙_ values compared to the CDG model (top panel of **Fig. 2B**). Together, these results indicate that positive priming can commonly occur with degraders exhibiting both copiotrophic and oligotrophic growth traits, though it is more facilitated by the former.

Meanwhile, CDG and ODG exhibited almost comparable growth over the entire range of the mixing fractions (bottom panels of **Fig. 2**). Owing to the inherent growth traits of copiotrophs and oligotrophs, the population level of CDG is low at pure complex OM (𝜈 = 0), while the population level of ODG is higher. However, at pure labile OM (𝜈 = 1), their population levels are comparable because, in the absence of complex OM to mine for additional labile OM, only the addition of exogenous labile OM remains for growth. Similar to the trend of the average 𝑃_𝑟𝑒𝑙_, the population levels of CDG and ODG significantly increased around 𝜈 ≈ 0.10, plateaued between 𝜈 ≈ 0.10 and 0.97, and showed no appreciable growth as 𝜈 approaches 1. Despite these general similarities, the average 𝑃_𝑟𝑒𝑙_ values and population densities showed major differences in the shapes of their profiles. In contrast with the average 𝑃_𝑟𝑒𝑙_ profiles showing a slight increase in the plateau, biomass concentration profiles showed a slight decline. This resulted in different optimal mixing fractions for maximizing the average 𝑃_𝑟𝑒𝑙_ and biomass concentrations, i.e., the former peaked around 𝜈 ≈ 0.97, while the latter reached its maximum around 𝜈 ≈ 0.1.

### 3.2 The impact of microbial interactions on priming is dependent on growth traits

We extended our analysis to the CDG–OND and ODG–CND mixed consortia (**Fig. 1D**) to examine how trait interactions affect priming. We hypothesized that non-degraders would suppress positive priming by weakening positive feedback loops and limiting the growth of degraders. Our results provide evidence that is only partially consistent with this hypothesis. We observed the opposite of what we expected in the CDG-OND consortium, whereby positive priming was promoted by OND microbes. This is indicated by the higher average 𝑃_𝑟𝑒𝑙_ values (top panel of **Fig. 3A**) compared to the CDG model (top panel of **Fig. 2A**). In contrast, and consistent with our hypothesis, we observed the frequent occurrence of negative priming in the ODG-CND consortium, resulting in lower average 𝑃_𝑟𝑒𝑙_ values (top panel of **Fig. 3B**), compared to the ODG model (top panel of **Fig. 2B**).

**Figure 3.**
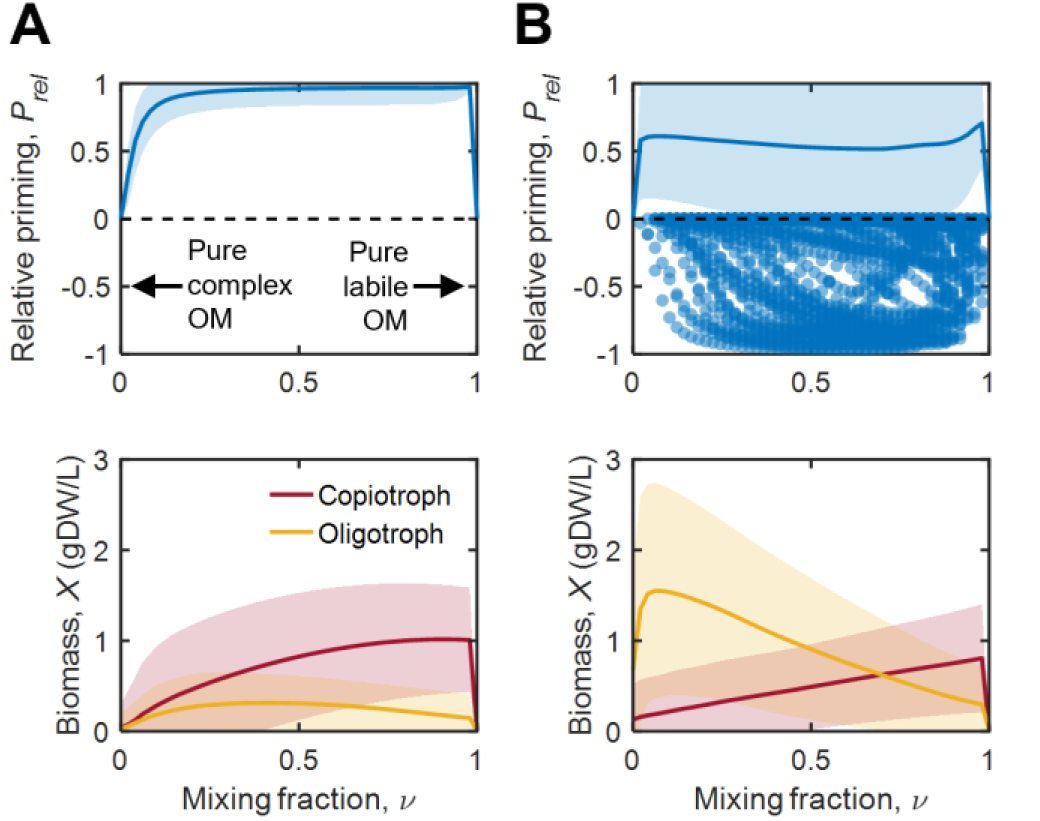
Overall relative priming effects and microbial population levels at various mixing compositions of complex and exogenous labile OM, evaluated for mixed cultures of **(A)** copiotrophic degrader and oligotrophic non-degrader, and **(B)** oligotrophic degrader and copiotrophic non-degrader. The lines and shaded regions are averaged results and standard deviations of Monte Carlo simulations, respectively, while the markers are individual runs with an overall negative priming effect.

In the CDG-OND consortium, the population level of degraders was consistently higher than that of non-degraders across the entire range of 𝜈 (bottom panel of **Fig. 3A**). This difference became more prominent at higher mixing fractions, conditions under which CDG can have growth advantages over OND. In contrast, the population densities of degraders and non-degraders showed opposing trends in the ODG-CND consortium along the mixing fraction (bottom panel of **Fig. 3B**). While ODG grew more effectively than CND at low values of mixing fractions, the dominance of degraders did not lead to increased priming due to the intrinsic nature of ODG, which is less effective at promoting positive priming. Moreover, the level of positive priming was nearly the same at both higher and lower values of 𝜈, where CND grows more effectively than ODG in the former.

### 3.3 Temporal dynamics of priming effects are shaped by microbial growth traits and interactions

In the previous two sections, we quantified overall relative priming over a specified reaction period, determined by an end time point, 𝑡_∞ (see **Methods**). The overall relative priming represents the cumulative sum of instantaneous priming effects at each time point over the given period. To understand how priming evolves over time, we also examined the temporal dynamics of instantaneous relative priming (𝑃_𝑖𝑛𝑠_) among the four focal priming models: CDG only (**Fig. 4A**), ODG only (**Fig. 4B**), CDG-OND consortium (**Fig. 4C**), and ODG-CND consortium (**Fig. 4D**), with three mixing fractions (𝜈 = 0.1, 0.5, and 0.9). In all cases, the average 𝑃_𝑖𝑛𝑠_ exhibits a unimodal trend, i.e., gradually increasing and then decreasing. Furthermore, the occurrences of negative average 𝑃_𝑖𝑛𝑠_ were more prevalent when the proportion of exogenous labile OM in the mix was high, especially during the later phases when positive priming weakened.

**Figure 4.**
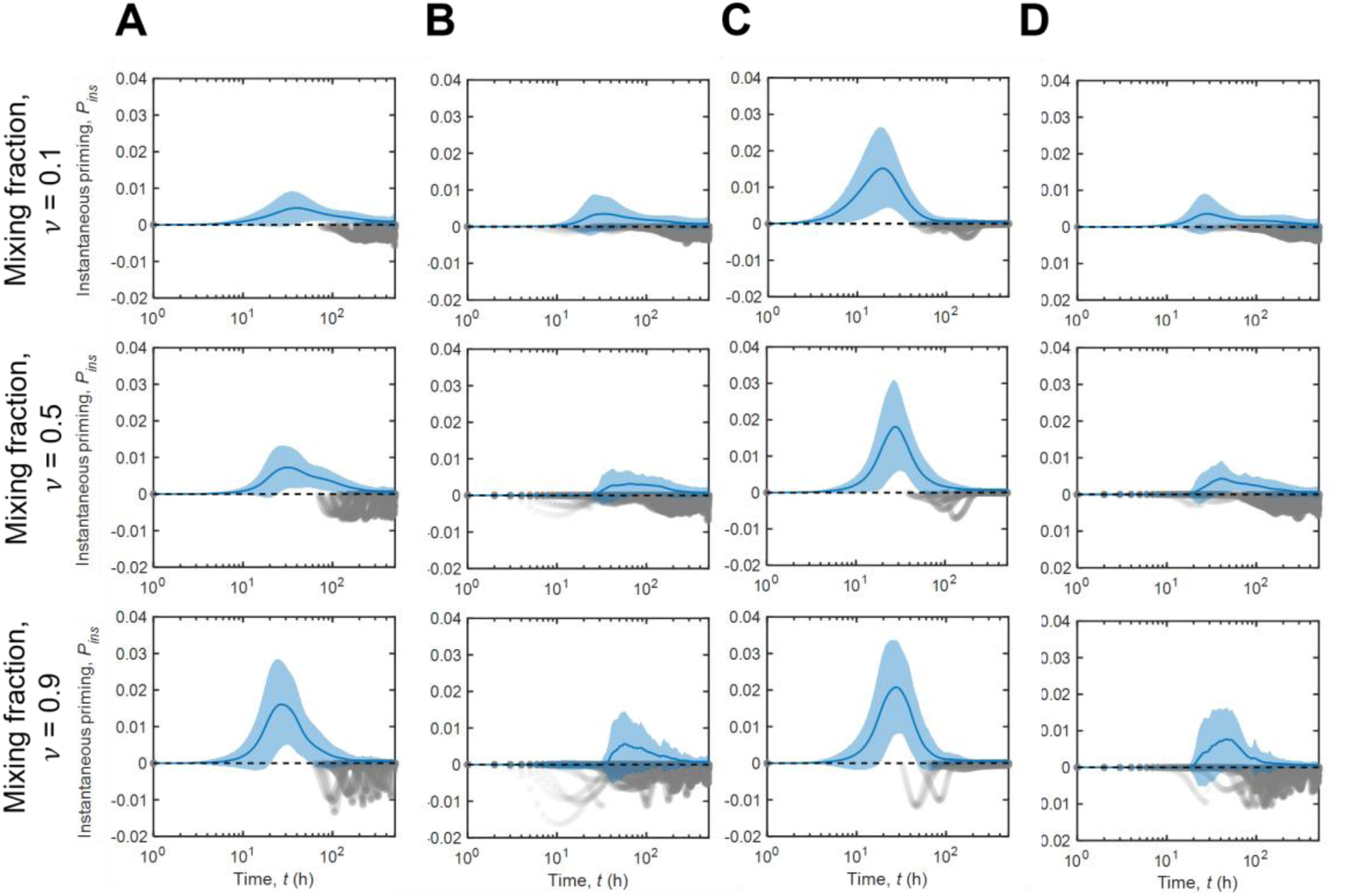
Instantaneous relative priming effects at low (𝜈 = 0.1), medium (𝜈 = 0.5), and high (𝜈 = 0.9) mixing compositions of exogenous labile OM with complex OM, evaluated for single functional group models of **(A)** copiotrophic degraders, **(B)** oligotrophic degraders, as well as binary consortia of **(C)** copiotrophic degrader and oligotrophic non-degrader, and **(D)** oligotrophic degrader and copiotrophic non-degrader. The lines and blue-shaded regions are averaged results and standard deviations of Monte Carlo simulations, respectively, while the gray-shaded markers indicate data points over time with negative instantaneous relative priming effects in individual runs.

Dynamics of priming were primarily dependent on the growth traits of degraders involved in the process. For example, the timing of maximum positive priming was consistent across mixing fractions in the CDG and CDG-OND models, where degraders exhibit copiotrophic growth traits (**Figs. 4A** and **4C**). In contrast, in the ODG and ODG-CND models, where degraders have oligotrophic growth traits, the peak times of positive priming shifted to later times with higher mixing fractions (**Figs. 4B** and **4D**). This is because CDGs grew more when more exogenous labile OM was added to the system, leading to quicker degradation of complex OM, compared to ODG whose dynamics were slower under the same conditions. Positive priming was more prominent when CDG was co-growing with OND (**Fig. 4C**), compared to the case with CDG alone (**Fig. 4A**). However, those synergies between degraders and non-degraders were not observed from the systems with ODG (**Fig. 4B** vs. **4D**), in line with our observations of the overall relative priming effects in **Fig 3**.

## 4 Discussion

Despite extensive research, the key mechanisms and factors governing priming remain elusive due to several challenges and complexities, including (1) the influence of numerous interacting biotic and abiotic factors, (2) difficulties in modeling the dynamics of microbial communities composed of multiple microbial functional groups with distinct growth traits, and (3) uncertain governing factors associated with microbial OM decomposition. A new modeling framework developed in this work enables systematic investigation in a large parameter space to reveal processes governing priming. With a focus on the impacts of key chemical and biological parameters (such as mixing fractions of complex and labile OM, microbial growth traits and interactions), we developed dynamic models of OM decomposition involving single functional microbial groups and consortia with distinct growth traits (copiotrophs and oligotrophs). We theorized the feedback loops of OM decomposition regulated by microorganisms to be a fundamental mechanism controlling the priming effects, which were incorporated into models using a cybernetic approach. This new microbial and OM decomposition model revealed several aspects of priming as detailed in the following sections.

### 4.1 Prevalence of positive priming and sporadic negative priming effects

Our modeling study shows that positive priming is prevalent in all test cases, and a small (less than 10%) addition of exogenous labile OM is sufficient to cause significant positive priming effects (top panels of **Figs. 2 and 3**). This is because a small amount of labile OM elicits notable microbial responses that lead to positive priming. This can be intuitively explained using the cybernetic perspective, as the microbial regulatory feedback loop shown in **Fig. 1B** is readily activated by degraders with either copiotrophic or oligotrophic growth traits by the addition of more bioavailable labile OM. The degraders gain a quick increase of energy from the assimilation of exogenous labile OM, allowing them to rapidly synthesize exoenzymes that degrade complex OM. This process produces more labile OM and perpetuates the feedback loop. This positive loop continues until the benefits of exoenzyme synthesis no longer outweigh the costs due to the significant depletion of complex OM. From an alternative perspective, investing in OM decomposition can be beneficial to degraders even in the absence of exogenous labile OM, as future growth may be supported by labile products generated during degradation. However, there may be insufficient available energy to initially synthesize the exoenzymes required for this process. The presence of exogenous labile OM removes this energy limitation, which is the fundamental working principle of positive priming.

On the other hand, negative priming is sporadic and therefore not observed in averaged results, but it is observed in individual runs of Monte Carlo simulations (top panels of **Figs. 2** and **3**, and **Fig. 4**). The conditions for negative priming often require more complex and less common scenarios such as when microbes preferentially use labile OM to immediately support cell growth rather than synthesizing exoenzymes to degrade complex OM to produce more labile OM for future growth. As we operate on the basis that degraders regulate their metabolism with a focus on returns over a finite future time horizon achieved using cybernetic modeling based on optimal control systems (see **Methods**), microbes can anticipate future returns (Young and Ramkrishna, 2007), especially when complex substrates are involved. Therefore, from the cybernetic perspective, negative priming can only manifest when the cost of exoenzyme synthesis to degrade complex OM outweighs the future returns in terms of cell growth from the assimilation of labile OM from degraded complex OM. As previously mentioned, the addition of labile OM almost always removes the energy limitation for exoenzyme synthesis, making negative priming less common than positive priming. In negative priming, degraders lack the incentive to degrade complex OM and instead focus on consuming the available labile OM to build biomass.

Although negative microbial priming is relatively uncommon, it can play a vital role in preserving soil carbon stocks and promoting carbon sequestration (Guenet et al., 2018; Liang et al., 2023). While positive priming enhances energy and nutrient exchange between microbial communities and the plant rhizosphere, excessive positive priming can contribute to the long-term depletion of soil carbon (Liang et al., 2017). Microbial priming, whether positive or negative, has significant implications for the global carbon cycle (Liang et al., 2017), and our study suggests that microbial regulatory factors must be considered in any efforts to predict or control future soil carbon stocks.

### 4.2 Prevalence of negative priming in temporal dynamics

Instantaneous relative priming effects exhibit highly non-linear dynamics over time and are more likely to cross into negative priming compared to the overall relative priming measure (**Fig. 4**). Temporal patterns of priming were unimodal because degraders immediately gain energy from labile OM, which accelerates degradation of complex OM, resulting in a peak of positive priming. This leads to the accumulation of labile OM. Microbes then allocate more energy towards consuming this labile OM in later phases, leading to a decrease in positive priming and the observed negative priming effect. If the test cases in this study are evaluated over longer timeframes, more negative priming may be observed. This highlights a need for greater focus on the temporal dynamics of priming because initial positive priming can evolve into negative priming.

Several studies provide further support for a stronger emphasis on temporal dynamics of priming. For example, Zhou et al. (2021) showed that priming responses are highly sensitive to dynamic changes in environmental conditions, which significantly influenced priming variability. Specifically, Zhang et al. (2017) demonstrated that the priming effect is modulated by incubation time, with distinct temporal phases observed, such as brief instances of initial negative priming followed by positive priming, and a stabilized phase later on. Several studies have reported that initial negative priming could result from OM consumption being dominated by the labile fraction, as microorganisms preferentially consume labile OM over complex OM (Kuzyakov and Bol, 2006; Khan et al., 2007; Guenet et al., 2010). As alluded to earlier, this behavior potentially reflects conditions in which microbes allocate labile OM directly to biomass production rather than investing in exoenzyme synthesis to degrade complex OM, particularly when the cost of exoenzyme synthesis outweighs the returns from complex OM degradation. Conversely, other studies have shown that the fast growth of copiotrophs that consume labile OM was observed to cause a rapid increase in mineralization of complex soil OM (Nicolardot et al., 2007; Pascault et al., 2013; Fu et al., 2022), potentially resulting in heightened positive priming. Mineralization of less complex OM by oligotrophs has also been reported (Blagodatskaya et al., 2009; Fang et al., 2018), which may lead to reduced positive priming. These observed patterns resemble outcomes from our Monte Carlo simulation runs and are consistent with the unimodal dynamics observed in **Fig. 4**. These reports also support our model results showing that the peak in positive priming occurs later with ODGs than with CDGs, which peak earlier. In line with the reasoning above, Soong et al. (2020) emphasized the need to account for temporal variability in environmental conditions to better capture how organisms utilize soil carbon and nutrients, ultimately leading to more representative ecosystem models.

### 4.3 The impacts of microbial growth traits on positive and negative priming

Despite their distinct growth traits, CDG and ODG both use labile OM for energy, enabling them to produce exoenzymes that degrade complex OM into labile OM for further energy gains. However, due to the preference for resource-rich conditions, the positive regulatory feedback loop is strongly activated when CDG are present. This results in CDG causing more positive priming than ODG (top panels of **Figs. 2** and **3**).

Conversely, ODG led to more negative priming when alone or paired with CND, especially at higher labile OM loading (top panels of **Figs. 2B and 3B**). Oligotrophs are adapted to thriving in resource-poor conditions, so when confronted with high amounts of labile OM, they encounter more than they can consume. In this scenario, it is not beneficial for them to generate additional labile OM through the costly process of synthesizing exoenzymes. Instead, they focus on consuming the available labile OM for growth rather than secreting exoenzymes for complex OM degradation. When ODG are paired with CND, the extent and frequency of negative priming significantly increased (top panel of **Fig. 3B**). This occurs because CND rapidly consumes the labile OM, leaving insufficient energy for ODG to continue degrading complex OM.

On the contrary, the presence of OND enhanced the extent of positive priming by CDG (top panel of **Fig. 3A**). This is also similarly observed in instantaneous relative priming effects (**Fig. 4C**). Oligotrophs can deplete limiting nutrients (i.e., labile OM in this context) to concentrations well below the threshold required for copiotrophs to persist following the R* theory (Couso et al., 2023), where R* indicates the minimum resource concentration that allows a population to maintain itself at equilibrium. Consequently, the addition of labile OM represents a larger perturbation in the binary CDG-OND system than in the single CDG system, thereby producing a stronger positive priming response in the former.

In agreement with our findings, Yang et al. (2023) inferred that soil basal respiration rates are largely driven from copiotrophs through the utilization of labile carbon sources, explaining the strong positive priming effects in our models involving CDGs. On the other hand, Fu et al. (2022) correlated copiotrophs with negative priming effects, implying that they mainly consume labile OM, thereby hindering the degradation of native complex soil OM, which could be explained by our models with CNDs. Furthermore, Fu et al. (2022) also reinforced our model findings, showing that the microbial community composition clearly exhibits distinct successions during OM decomposition, leading to distinct priming patterns. However, while the studies above associate the variability in priming with the growth traits of microbial groups, they do not distinguish between degraders and non-degraders or consider the interactions between microbial groups. Our model does examine these factors, highlighting its value in providing insights across complex scenarios.

### 4.4 Microbial population dynamics and emergent priming responses

With regards to microbial population levels, CDG and ODG exhibit almost comparable growth when considered independently (bottom panels of **Fig. 2**). Conversely, in binary consortia, oligotrophs attain higher population levels than copiotrophs at low mixing fractions, and vice versa at high mixing fractions, as expected based on their growth traits (bottom panel of **Fig. 3B**). The only exception is when CDG is paired with OND, where the former dominates across all mixing fractions (**Fig. 3A**). This dominance occurs because the uptake of rich labile OM in OND is slower, preventing it from effectively capitalizing on the efforts of copiotrophic degraders in producing labile OM. Moreover, the population of OND stabilizes and ODG declines with increasing levels of exogenous labile OM (bottom panels of **Figs. 2** and **3**). This is because ODG is at a greater disadvantage in a labile OM-rich environment compared to OND because although both are unable to competitively utilize the surplus labile OM, ODG must continue synthesizing exoenzymes to maintain the labile OM level in the environment, incurring additional metabolic costs.

We also found that the priming magnitudes and population levels of microbial groups were not synchronized. This indicates that although one of the major factors driving microbial priming is biomass concentration, there are other influential factors. As degraders must expend significant resources to degrade complex OM to induce maximum positive priming, this leaves relatively less energy available for growth, resulting in lower population levels. This suggests that the microbial group exerting the strongest influence on priming (i.e., degraders) does not have to be the most abundant within the community. Because priming is a shared and robust feature across diverse microbial systems and environments, distinct community compositions can produce comparable net priming effects. Thus, priming patterns are better understood as emerging from community-level interactions rather than being tied to a single microbial group.

### 4.5 A small addition of exogenous labile OM triggers significant positive priming

One of the common features of priming that we identified in our work, consistent across all test cases, is that significant positive priming was triggered by the addition of small amounts of labile OM (less than about 10%), beyond which no further significant changes were observed. The variation in priming levels with increasing exogenous labile OM, along with the presence of a maximum positive priming level, indicates a threshold effect. When the addition of exogenous labile OM exceeds a certain threshold, there is more labile OM than degraders can consume. As a result, the system reaches a saturation point where degraders cannot further increase the exoenzyme synthesis rate, leading to no additional improvement in positive priming, even with more labile OM. Although this threshold may vary slightly between copiotrophs and oligotrophs, it is generally less than around 10% labile OM, depending on the fixed model parameters in this study. In addition, beyond the threshold where there is a surplus of labile OM, the motivation to further increase OM decomposition diminishes. As a result, microbes maintain their existing course of action, using the available labile OM to sustain exoenzyme synthesis and maintain the level of labile OM in the system through the degradation process.

Similar findings were reported by Stegen et al. (2018), where the addition of up to only 10% labile OM from groundwater to complex OM from river water resulted in a significant increase in overall OM oxidation in the mixture. However, no further increases in oxidation rate were observed beyond this 10% groundwater threshold. The increase in OM oxidation was also not significant well below 10% groundwater because the concentration of OM was so low that the energy acquired from the assimilation of labile OM was insufficient to offset the energetic costs required to oxidize it (Arrieta et al., 2015). Similarly, Guenet et al. (2010) suggested that the priming effect in soil does not vary linearly with the addition of labile OM, instead, the amount of labile OM only partially controls priming, which functions as a saturating response. This closely resembles the observation in our study, where significant positive priming plateaus beyond a certain threshold. Likewise, Zhou et al. (2021) observed that frequently adding smaller amounts of labile OM resulted in a greater priming response in soil than adding larger amounts less frequently. Lastly, one of the established mechanisms of positive priming is the acceleration of internal microbial metabolism by *trace amounts* of substrates, which leads to an immediate increase in microbial respiratory activity in soil (Blagodatsky et al., 2010; Bernard et al., 2022), consistent with our findings.

### 4.6 Conclusions

Priming effects can be complex due to the combined influence of numerous chemical and biological factors. To account for these complexities, we developed a modeling framework to systematically investigate the effects of mixing complex and labile OM. The framework further evaluates how microbial growth traits and interactions influence priming through regulatory mechanisms that govern complex OM decomposition. As a result, our study offers a mechanistic foundation for understanding microbial priming effects in natural ecosystems. Our results in combination with previous studies suggest that although the priming effect may occur in diverse environments such as hyporheic zones, soils, or other ecological contexts, some of its core features remain transferable across systems because the process is ultimately driven by microbial behavior that operates under common regulatory principles. Notably, the identification of a critical threshold, where a minimal addition of labile organic matter can trigger significant positive priming, highlights the sensitivity of microbial communities to environmental changes and the potential for small perturbations to have significant impacts on biogeochemical cycles. Moreover, microbial growth traits and interactions further influence the magnitude and direction of priming in multiple ways. For instance, our study revealed that copiotrophic degraders are generally associated with significant positive priming, while oligotrophic degraders and/or copiotrophic non-degraders are linked to sporadic occurrences of negative priming. These insights have broad implications for ecological modeling, biogeochemical research, and environmental management.

Some of our findings may be limited to the model conditions used in this study (e.g., fixed parameter values and constraints for randomized parameters representing copiotroph and oligotroph growth traits, complex OM properties, and complex OM degradation kinetics). There is potential for other significant observations to emerge when the model conditions are expanded to encompass a broader range of settings. For example, we have not considered the effects of steric hindrances of OM protected within the soil or mineral aggregates (Marschner and Kalbitz, 2003; Schmidt et al., 2011; Lavallee et al., 2020), or the mass transport of OM under non-uniform conditions. It is also important to consider the impact of OM stoichiometry and thermodynamic favorability on microbial processes and their implications for priming effects (Graham and Hofmockel, 2022). Existing literature highlights how variations in OM composition, especially C/N ratios, can profoundly influence microbial activity (Ahamed et al., 2023) and the dynamics of priming (Fontaine et al., 2011; Abramoff et al., 2017; Hicks et al., 2019). Furthermore, the availability or limitation of nutrients, such as nitrogen, can also influence priming responses (Chen et al., 2014; Fu et al., 2022).

Future research should focus on integrating theoretical models with empirical data to test the hypotheses generated from this work, while also incorporating other major governing factors highlighted above. Doing so will further refine our understanding of priming mechanisms and improve predictions of ecosystem responses to environmental perturbations. We expect that the key concepts proposed in our framework will contribute significantly to the advancement of predictive ecological models aimed at understanding elemental cycling in natural ecosystems.

## Author Contributions

All authors contributed to the design of the research. FA and H-SS developed the computational framework for exploring priming effects and drafted the article. FA developed numerical codes and performed computational simulations. The manuscript was reviewed, edited, and approved by all authors.

