## Supplementary material for "Modeling Microbial Regulatory Feedback in Organic Matter Decomposition Identifies Copiotrophic Traits as Key Drivers of Positive Priming": Supp text, figures S1 and S2, tables S1 and S2

### SUPPLEMENTARY TEXT

#### Growth Traits of Copiotrophs and Oligotrophs

In **Supplementary Fig. S1**, we showed the behavior of oligotrophs and copiotrophs at varying labile organic matter (OM) concentrations and respective population levels based on our fixed parameter values. While oligotrophs and copiotrophs can thrive in environments with less and more than 0.1 M of labile OM (**Supplementary Fig. S1A**), the microbial activities are also influenced by their respective population levels. The copiotrophs are more sensitive to changes in population levels, where they need to exceed the oligotroph population by more than three times under labile OM-limited conditions, and by 1.5 times under labile OM-rich conditions to equally compete for available labile OM (**Supplementary Fig. S1B**). Therefore, we set equal initial population levels for all microbial groups in all test cases in this study to give them equal footing in accessing labile OM (see **Methods**). However, the complex dynamics of complex OM degradation, simultaneous labile OM uptake, and microbial population accumulation can all influence the regulation strategy of microbes. Additionally, the degraders can transform the availability of labile OM for non-degraders depending on their efficiency in degrading complex OM. Therefore, it is imperative that we not only use the end conditions to characterize the overall priming effects but also consider the transient changes in priming effects over time. The overall and transient priming outcomes for various test cases are explored in the **Results** and **Discussion** section.

#### Priming Effects and Microbial Population Levels in Alternative Binary Consortia Configurations

We further explored additional binary consortia configurations, including ODG-OND, CDG-CND, and CDG-ODG, that were not covered in the main text. When both CDG and ODG are paired together, the extent of positive priming is not as high, and occurrences of negative priming have significantly increased compared to the single functional group model of CDG or the binary consortium of CDG and OND (cf. top panels of **Supplementary Fig. S2A** with **Fig. 2A** and **Fig. 3A**, respectively). This can be explained by the presence of two degraders potentially leading to the accumulation of labile OM in the system, resulting in a decline in the degradation of complex OM. As expected from their characteristic growth traits, ODG fare better in labile OM-poor conditions, while CDG thrive in labile OM-rich conditions (bottom panel of **Supplementary Fig. S2A**). Conversely, when degraders and non-degraders share similar growth traits, they compete equally for labile OM, resulting in less efficient triggering of positive priming effects, and increased chances of negative priming when compared with the single functional group models of the respective degraders (cf. top panels of **Supplementary Figs. S2B** and **S2C** with **Fig. 2**). Similarly, the combination of CDG and CND tends to have a higher frequency of negative priming compared to when CDG are paired with OND (cf. top panels of **Supplementary Fig. S2B** with **Fig. 3A**). This is because CND in the former are more adept at utilizing the labile OM produced by CDG, thereby reducing the potential of positive priming. Also, CDG and CND have comparable population levels as they both equally thrive and compete in labile OM-rich conditions (bottom panel of **Supplementary Fig. S2B**). However, the combination of ODG and OND shows fewer instances of negative priming compared to when ODG are paired with CND (cf. top panels of **Supplementary Fig. S2C** with **Fig. 3B**). This is because OND in the former are less effective in exploiting the degradation products produced by ODG.

### SUPPLEMENTARY FIGURES

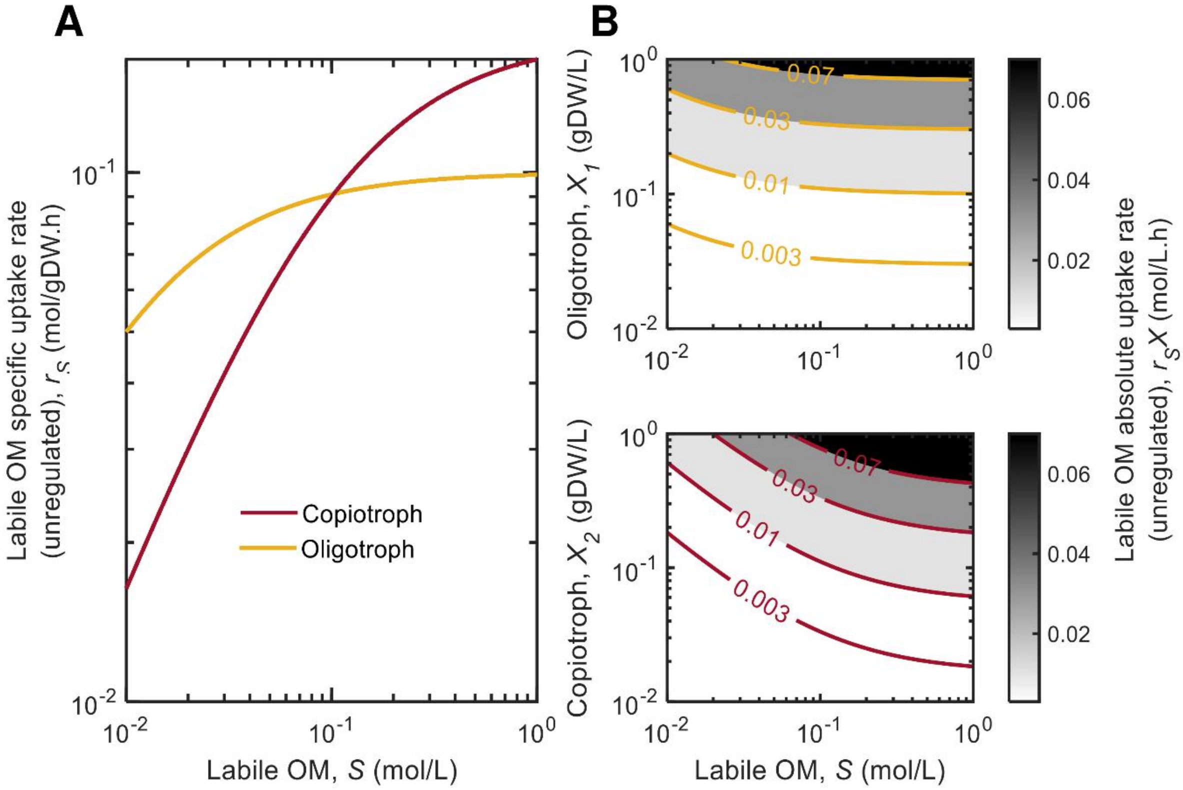


**Supplementary Fig. S1.** Differences in labile OM uptake kinetics between oligotrophs and copiotrophs visualized as **(A)** unregulated specific uptake rates with respect to only labile OM levels, and **(B)** unregulated absolute uptake rates with respect to both biomass and labile OM levels.

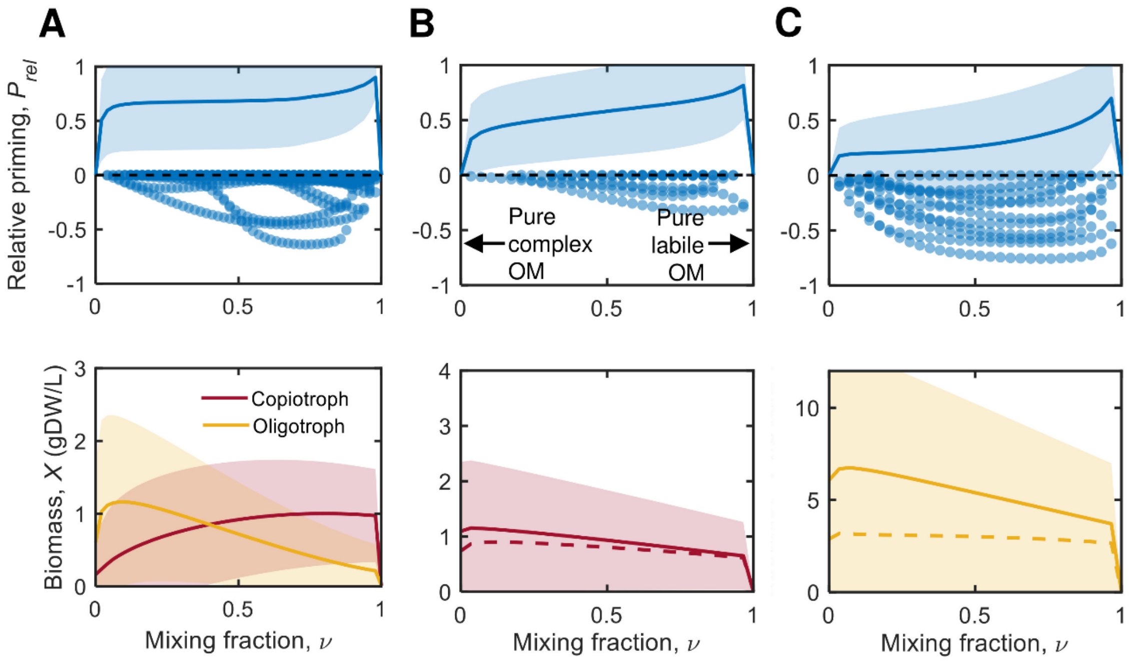


**Supplementary Fig S2.** Overall relative priming effects and microbial population levels at various mixing compositions of complex and exogenous labile OM, evaluated for binary consortia of **(A)** oligotrophic and copiotrophic degraders, **(B)** copiotrophic degraders and non-degraders, and **(C)** oligotrophic degraders and non-degraders. The lines and shaded regions are averaged results and standard deviations of Monte Carlo simulations, respectively, while the markers are individual runs with negative overall relative priming effects. In the population-level plots, the solid lines are degraders and dashed lines represent non-degraders.

### SUPPLEMENTARY TABLES

**Supplementary Table S1.** Model parameter values that were either fixed or randomized within specified bounds using uniform distributions in this study.

| Model parameter | Fixed [F] or Randomized [R] | Copiotrophs | Oligotrophs | Remarks |
|---|---|---|---|---|
| $Y_{X,i}$ | R | [0, 2] | [0, 2] | - |
| $k_{S,i}$ | F | 0.18 | 0.1 | Copiotrophs > oligotrophs |
| $K_{S,i}$ | F | 0.1 | 0.01 | Copiotrophs > oligotrophs |
| $\alpha_{E,i}$ | R | [0, 1E-4] | [0, 1E-4] | - |
| $\alpha_{e,i}$ | R | $= \alpha_{E,i}$ | $= \alpha_{E,i}$ | Assuming basal synthesis rates of exoenzymes and endoenzymes are identical for simplicity. |
| $Y_{E,i}$ | R | [0, 2] | [0, 2] | - |
| $Y_{e,i}$ | R | [0, 2] | [0, 2] | - |
| $\beta_{E,i}$ | F | 0.01 | 0.01 | - |
| $\beta_{e,i}$ | F | 0.01 | 0.01 | - |
| $K_{E,i}$ | R | [0, 0.1] | [0, 0.1] | - |
| $K_{e,i}$ | R | [0, 0.1] | [0, 0.1] | - |
| $X_i(0)$ | F | 0.01 | 0.01 | Made equal for all microbial groups for a fair comparison. |
| $Z(0)$ | F | Depending on the mixing fraction, $\nu$ | | $\begin{cases} \nu = 0: Z(0) = 1; S(0) = 0 \\ \nu = 1: Z(0) = 0, S(0) = 1 \end{cases}$ |
| $S(0)$ | F | | | |
| $k_Z$ | F | 0.01 | | Fixed to exclude the effects of variability in complex OM properties on priming. |
| $K_Z$ | F | 0.1 | | |
| $Y_S$ | F | 2 | | |

**Supplementary Table S2.** Summary of results for various priming models with different assortments of microbial groups exhibiting distinct growth traits. The levels of priming (low: $P_{rel} < 0.1$, moderate: $P_{rel} = 0.1 - 0.9$, high: $P_{rel} > 0.9$) and the percentage occurrences (runs) of different priming types (positive, negative, zero) in the Monte Carlo simulations are presented. Note: CDG – copiotrophic degrader; ODG – oligotrophic degrader; CND – copiotrophic non-degrader; OND – oligotrophic non-degrader; PP – positive priming; NP – negative priming; ZP – zero priming.

| | CDG | ODG |
|---|---|---|
| CDG | PP: High levels, 96.0%<br>NP: 0.0%<br>ZP: 4.0% | PP: Moderate to high levels, 94.6%<br>NP: Low to moderate levels, 1.4%<br>ZP: 4.0% |
| ODG | PP: Moderate to high levels, 94.6%<br>NP: Low to moderate levels, 1.4%<br>ZP: 4.0% | PP: Moderate to high levels, 95.7%<br>NP: Low to high levels, 0.3%<br>ZP: 4.0% |
| CND | PP: Low to moderate levels, 88.0%<br>NP: Low to moderate levels, 5.3%<br>ZP: 6.7% | PP: Moderate levels, 90.4%<br>NP: Low to high levels, 5.6%<br>ZP: 4.0% |
| OND | PP: High levels, 100.0% PP<br>NP: 0.0%<br>ZP: 0.0% | PP: Low to moderate levels, 82.9%<br>NP: Low to moderate levels, 10.4%<br>ZP: 6.7% |
